# Regulatory sites in the Mon1-Ccz1 complex control Rab5 to Rab7 transition and endosome maturation

**DOI:** 10.1101/2023.03.01.530579

**Authors:** Ann-Christin Borchers, Maren Janz, Jan-Hannes Schäfer, Arne Moeller, Daniel Kümmel, Achim Paululat, Christian Ungermann, Lars Langemeyer

## Abstract

Maturation from early to late endosomes depends on the exchange of their marker proteins Rab5 to Rab7. This requires Rab7 activation by its specific guanine nucleotide exchange factor (GEF) Mon1-Ccz1. Efficient GEF activity of this complex on membranes depends on Rab5, thus driving Rab-exchange on endosomes. However, molecular details on the role of Rab5 in Mon1-Ccz1 activation are unclear. Here we identify key features in Mon1 involved in GEF regulation. We show that the intrinsically disordered N-terminal domain of Mon1 autoinhibits Rab5-dependent GEF-activity on membranes. Consequently, Mon1 truncations result in higher GEF activity *in vitro*, and a shift from Rab5 to more Rab7 positive structures in *Drosophila* nephrocytes and yeast, suggesting faster endosomal maturation. Using modeling, we further identify a conserved Rab5 binding site in Mon1. Mutations impairing Rab5 interaction result in poor GEF activity on membranes and growth defects *in vivo*. Our analysis provides a framework to understand the mechanism of Rab-conversion and organelle maturation along the endomembrane system.

**Summary:** Transport of proteins via the endolysosomal pathway requires Rab5 on early endosomes, which is replaced by Rab7 on late endosomes. Here, we identify distinct regulatory sites in the Rab7 activator, the Mon1-Ccz1 complex, shedding light on the regulation of Rab5 to Rab7 transition.

## Introduction

All eukaryotic cells utilize endocytosis for homeostasis of their plasma membrane and exchange of materials with the cell’s surrounding. Endocytic cargo is packed into endocytic vesicles, which pinch off at the plasma membrane and fuse with early endosomes. Early endosomes fuse and mature into late endosomes, finally delivering endocytic cargo to the degradative lysosome. Two members of the Rab-subfamily of small GTPases, Rab5 and Rab7, are critical identity markers within the endolysosomal pathway (Borchers et al., 2021). Rab5 decorates early endosomal compartments. During maturation of early to late endosomes Rab5 is replaced by Rab7. This transition is critical for the functionality of each organelle, and a shift in this fine balance can lead to diseases like the Charcot-Marie-Tooth syndrome type 2 (Romano et al., 2021).

Rab-GTPases (Rabs) are guanine-nucleotide binding proteins with a weak intrinsic GTPase activity. Rabs are prenylated at their C-terminus to allow for membrane association and cycle between a soluble GDP- and a membrane-anchored GTP-bound state. Within the cytosol, the GDP dissociation inhibitor (GDI) binds to the GDP-bound form and covers its prenyl anchor. The region between GTPase domain and prenyl anchor, called hypervariable domain (HVD), is rather flexible. Rabs can dissociate from GDI and thereby sample membranes in the cell. If they encounter their cognate guanine nucleotide exchange factor (GEF) at their target organelle, GDP is exchanged for the more abundant GTP. In this active GTP-bound state, Rab-GTP stably associates with the membrane and can recruit and bind effector proteins, for instance tethers or motor proteins. Specific GTPase activating proteins (GAP) can bind to their respective Rab and catalyze the GTP-hydrolysis, thereby transferring the Rab back to its inactive state which can be extracted by GDI (Goody et al., 2017; Barr, 2013; Borchers et al., 2021).

On early endosomes, and possibly already on endocytic vesicles, Rab5 is activated by its GEF Rabex5. Active Rab5 recruits, among others, its effector Rabaptin. Rabaptin in turn interacts with Rabex5 resulting in a positive feedback-loop establishing Rab5-domains on endosomal membranes (Bezeljak et al., 2020; Cezanne et al., 2020). Other effectors of Rab5 are for instance the tethering complex CORVET and the phosphatidylinositol-3-phosphate (PI3P) kinase complex II (Kümmel and Ungermann, 2014; Tremel et al., 2021).

An additional Rab5 effector is the GEF for the late endosomal Rab7, the Mon1-Ccz1 complex (Kinchen and Ravichandran, 2010; Langemeyer et al., 2020; Cui et al., 2014; Singh et al., 2014), a heterodimer in yeast and a heterotrimer in metazoan cells (Wang et al., 2002; Vaites et al., 2018; Boomen et al., 2020; Dehnen et al., 2020). The third subunit, called RMC1/C18orf8 in humans or Bulli in *Drosophila*, is required for a functional endosomal system and autophagy in metazoan cells (Vaites et al., 2018; Boomen et al., 2020; Dehnen et al., 2020). However, GEF activity of the *Drosophila* Mon1-Ccz1 dimer and trimer is comparable, suggesting that the third subunit functions in other processes rather than regulating GEF activity (Dehnen et al., 2020; Langemeyer et al., 2020). Mon1-Ccz1 belongs to the family of Tri-Longin-GEFs, in which each subunit consists of three Longin-domains (LDs). This family includes the BLOC3 complex of Hps1 and Hps4, a GEF of Rab32 and 38 involved in the biogenesis of lysosome-related organelles (Gerondopoulos et al., 2012), and the CPLANE complex with its three subunits Fuzzy, Inturned, and Wdpcp, which also binds Rsg1 as a non-canonical Rab-like subunit (Langousis et al., 2022). Within Mon1-Ccz1, LD1 of both subunits forms the active site (Kiontke et al., 2017; Klink et al., 2022), whereas LD2 and LD3 of both proteins form a layer responsible for membrane association (Herrmann et al., 2023). Rab7 activation by Mon1-Ccz1 is necessary for fusion events of late endosomes and autophagosomes with the lysosome.

Transition from a Rab5-positive to a Rab7-positive domain on the verge from early to late endosomes has been termed Rab-cascade (Goody et al., 2017; Barr, 2013; Borchers et al., 2021; Hutagalung and Novick, 2011; Conte-Zerial et al., 2008). Mon1-Ccz1 is an effector of Rab5 and is recruited to Rab5-positive membranes, and subsequently activates Rab7 (Fig. 1A)(Kinchen and Ravichandran, 2010; Langemeyer et al., 2020; Cui et al., 2014). Using reconstitution, we showed before that presence of active Rab5 on membranes increases GEF-activity of Mon1-Ccz1 suggesting an activation of the GEF-complex on endosomal membranes on the verge to late endosomes (Langemeyer et al., 2020; Herrmann et al., 2023). However, it remains unclear how binding to Rab5 activates Mon1-Ccz1. It is also unresolved, how a sharp spatiotemporal transition between Rab5- and Rab7-domains in the endolysosomal pathway is achieved, and why Mon1-Ccz1 is not activated by Rab5 on early endosomal structures.

**Figure 1.**
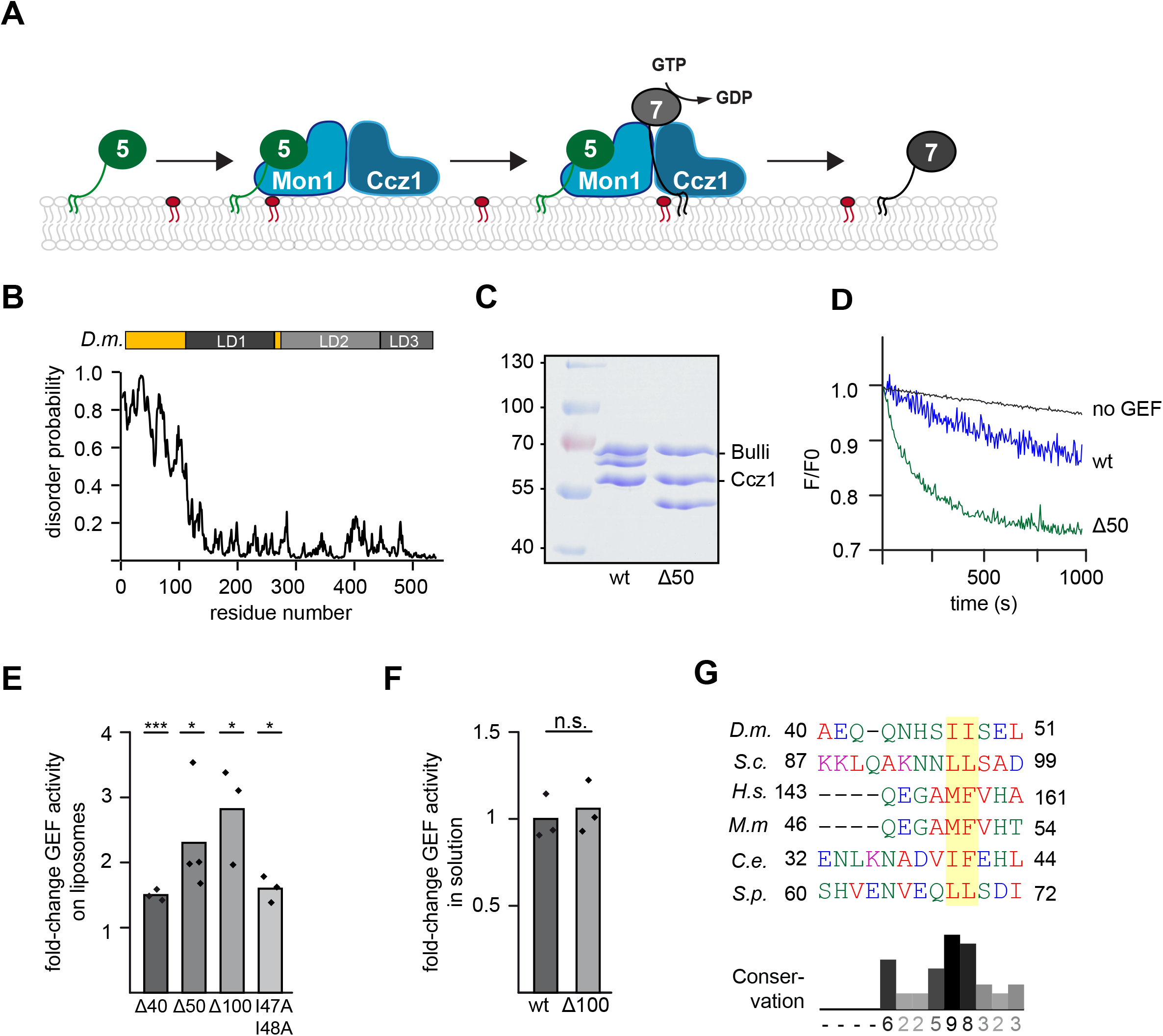
The disordered N-terminal region of *Drosophila* Mon1 regulates Mon1-Ccz1 GEF activity. (**A**) Overview of Rab5-Rab7 cascade on endosomes. Active Rab5 (green) recruits the Mon1-Ccz1 GEF complex (blue) to PI3P positive (red) endosomal membranes and promotes Rab7 (grey) recruitment and activation. For details see text. (**B**) The N-terminal region of *D.m*. Mon1 is disordered. Disorder probability of each residue of *D.m*. Mon1 was determined using IUPred2A web interface (Erdős and Dosztányi, 2020; Mészáros et al., 2018). Longin domains (LD) 1-3 are depicted in grey shades. Values >0.5 are considered as disordered. (**C**) Truncation of Mon1 does not affect complex stability of the trimeric Mon1^Δ1-50^-Ccz1-Bulli complex (Δ50) compared to wild-type complex (wt). GEF complexes were purified as described in the method section and analyzed by SDS-PAGE and Coomassie staining. (**D**) On liposome GEF assay of Mon1^wt^ and Mon1^Δ1-50^ containing trimer. Liposomes were preloaded with 150 nM prenylated Rab5 in the presence of 200 μM GTP and 1.5 mM EDTA. Nucleotide was stabilized using 3 mM MgCl_2_. 250 nM Mant-GDP loaded Rab7:GDI were added and nucleotide exchange was triggered by adding 6.25 nM wild-type (blue) or mutant (green) GEF complex. Decrease in fluorescence was measured over time and normalized to fluorescence prior to GEF addition. (**E**) Comparison of fold-change in GEF activity of various Mon1 truncations and mutants. GEF assays were performed as in (**D**) and k_obs_ of each curve was determined as described in the method section. k_obs_ values of mutants were normalized to the corresponding wild-type value. Bar graphs represent average foldchange and dots represent individual changes from at least three experiments. (P-value * p<0.05, ** p<0.01, using two sample t-test). For kinetic constants of all GEF complexes, see Table S3. (**F**) GEF activity of wildtype and Mon1^Δ1-100^ containing trimer in solution. 2 μM non-prenylated Rab were incubated with increasing amounts of GEF. After baseline stabilization, nucleotide release was triggered by adding 0.1 mM GTP final. For details of data fitting and statistics see method section. The k_cat_/K_M_ (M^-1^s^-1^) value for Mon1^Δ1-100^ containing trimer was normalized to the value of Mon1^wt^. Bar graphs represent average fold-change, and dots represent individual changes from three experiments. (P-value n.s. using two sample t-test). (**G**) Multiple sequence alignment of a hydrophobic patch in Mon1. The N-terminal regions of Mon1 from *Drosophila melanogaster (D.m.), Saccharomyces cerevisiae (S.c.), Homo sapiens (H.s.), Mus musculus (M.m.), Caenorhabditis elegans (C.e.*) and *Schizosaccharomyces pombe (S.p.*) were aligned using Clustal omega web interface (Sievers et al., 2011; Goujon et al., 2010). Hydrophobic patch is marked by a yellow box. Conservation was determined using Jalview (Waterhouse et al., 2009).

To address these questions, we investigated here a possible regulatory function of the mainly intrinsically disordered Mon1 N-terminal region. We show that this domain autoinhibits Mon1-Ccz1 and further identify a conserved Rab5 binding site in yeast and *Drosophila* Mon1 by structural modeling. Our results provide a framework on the regulation of GEFs, which thus control Rab transitions during organelle maturation.

## Results

### Mon1-Ccz1 activity is autoinhibited

We showed before that activity of the Rab7-GEF Mon1-Ccz1-Bulli is stimulated by Rab5 in a reconstituted system (Langemeyer et al., 2020; Dehnen et al., 2020), suggesting an order of events that promotes endosomal maturation (Fig. 1A)(Borchers et al., 2021). In the past, we determined the structure of the dimeric Mon1-Ccz1 complex and identified the overall organization of the complex and a unique mechanism to drive nucleotide exchange of Rab7 (Kiontke et al., 2017; Klink et al., 2022). However, in both structures the N-terminal domain of Mon1 was not resolved. This missing part is predicted to be disordered in *Drosophila melanogaster* (*D.m.*) (Fig. 1B), *Saccharomyces cerevisiae* (*S.c.*) and human Mon1 (Fig. S1A,B). This feature is conserved and unique among TLD subunits, suggesting a functional role of this region. We generated truncations of the N-terminal part and analyzed purified Mon1-Ccz1-Bulli complexes *in vitro*. All complexes were purified as a trimer (Fig. 1C and Fig. S1C).

To test for activity, we used an established *in vitro* GEF-assay (Langemeyer et al., 2020). In brief, Mon1-Ccz1-Bulli activity was determined by following the release of a fluorescently labeled GDP (Mant-GDP) from a preloaded Rab7-GDI complex in the presence of unlabeled GTP upon GEF addition. To mimic the membrane environment of the cell, this assay was conducted in the presence of liposomes, which carried prenylated Rab5-GTP that is required for efficient Rab7 GEF activity (Langemeyer et al., 2020). We then tested the effect of Mon1-truncations on GEF activity. Compared to the wild-type complex, N-terminal truncations of the first 40, 50 or 100 residues resulted in a 1.5-3.5-fold increase in GEF-activity (Fig. 1D,E and Table S3). This suggested that the N-terminal unstructured region inhibits Mon1-Ccz1. Surprisingly, GEF activity of the most active Mon1-Ccz1 mutant was like wildtype in solution using non-prenylated, soluble Ypt7 as a substrate (Fig. 1F). Closer inspection of the region between residues 40-50 of *D.m*. Mon1 revealed a conserved hydrophobic motif in this region (Fig. 1G), A conservative I47, 48A mutant also resulted in an increase in Rab5-dependent Mon1-Ccz1-Bulli activity compared to wildtype (Fig. 1E), indicating that this hydrophobic patch regulates the availability of Mon1 and thus controls GEF complex activity.

### The regulatory role of the Mon1 N-terminal region is conserved

We next asked if we could observe an effect of the Mon1 N-terminal truncation in *Drosophila in vivo*. In previous work, we analyzed the effect of Bulli deletions on the morphology and function of nephrocytes. *Drosophila* nephrocytes share histological and functional similarities with mammalian kidney podocytes, for example, a filtration apparatus and a complex endocytic system (Dehnen et al., 2020; Psathaki et al., 2018). As the *Drosophila* GEF complex carrying Mon1^Δ100^ had higher GEF activity than the wildtype (Fig. 1E), we expected that expression of Mon1^Δ100^ in *Drosophila* nephrocytes would generate a dominant effect on the morphology of the endocytic system. Therefore, a *Drosophila* line was generated expressing Mon1^Δ100^, which was verified by Western Blot analysis (Fig. S2A). As Bulli mutants had a strongly impaired ultrastructural morphology (Dehnen et al., 2020; Psathaki et al., 2018), we initially analyzed nephrocytes by electron microscopy. The expression of Mon1^Δ100^ had, however, little visible consequences on the histology of nephrocytes. Ultrastructural analyses revealed similar slit diaphragms and a normal labyrinth channel system, representing the endocytic compartment. The slit diaphragms serve as a filtration barrier. We also identified coated vesicles, early endosomes, and other typical organelles of nephrocytes (Fig. S2B). The expression of Mon1^Δ100^ also did not affect the endocytic activity of nephrocytes, as tested by uptake assays with FITC albumin (Fig. S2C,D). These observations were not unexpected, as the Mon1^Δ100^ supports Rab7 recruitment, and subtle defects in endolysosomal transport may not become apparent by these assays. However, the allele may nevertheless change the apparent transition from early to late endosomes.

To determine possible effects on the Rab transition in the endosomal system, we stained nephrocytes expressing Mon1^Δ100^ for Rab5 and Rab7, and observed a clear increase in colocalization of both Rabs, indicating that they remain longer on the same endosomal compartments, possibly due to premature activation of Rab7 on early endosomes (Fig. 2A,B). Moreover, Rab5 localized to more and smaller vesicular compartments within Mon1^Δ100^ expressing nephrocytes than in control cells. We thus conclude that the Mon1^Δ100^ allele causes a shift in the balance from early to late endosomes and thus enhance the Rab5 to Rab7 transition in nephrocytes, while still allowing for endocytic transport.

**Figure 2.**
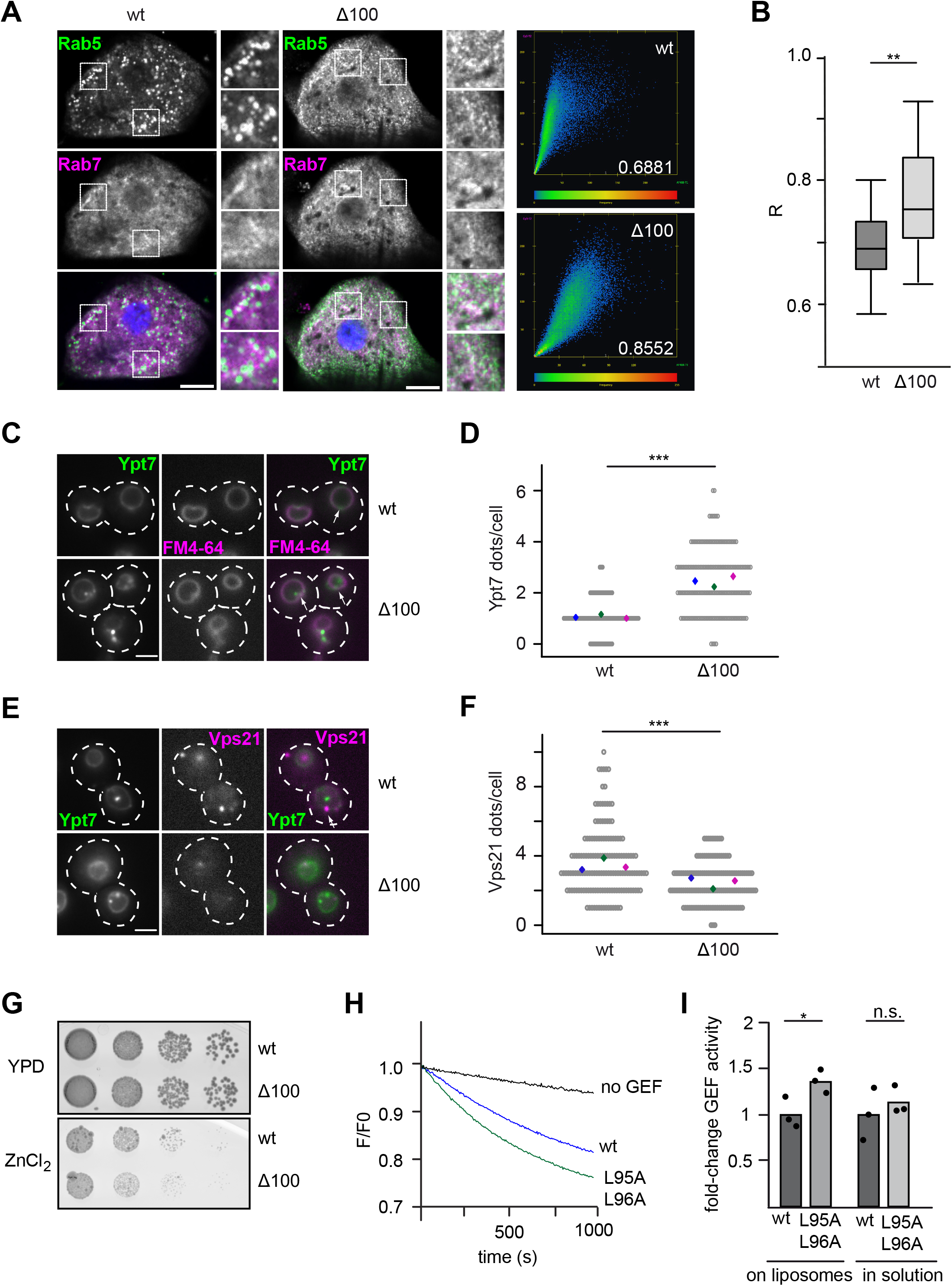
Loss of the disordered Mon1 N-terminal region affects localization of endosomal Rab GTPases and Rab5-dependent GEF activity. **(A)** Localization of endogenous Rab5 and Rab7 in *Drosophila* nephrocytes from 3^rd^ instar larvae expressing wildtype or Mon1^Δ100^ under the control of the *handC*-GAL4 driver. Antibodies against Rab5 and Rab7 were used. Optical sections show the distribution of Rab5 and Rab7 in detail. Size bar: 10 μm. Corresponding scatter plots for nephrocytes showing the Pearson correlation coefficient R. (**B**) Quantification of the Pearson correlation coefficient R calculated from 17 cells from three animals for wild-type Mon1 and 12 cells from four animals for Mon1^Δ100^. (P-value **<0.01 using two sample t-test). **(C-F)**Truncation of the Mon1 N-terminal affects Ypt7 and Vps21 localization. **(C)**Localization of mNEON-Ypt7 under its endogenous promotor in yeast cells in the presence of endogenously expressed wildtype or Mon1^Δ100^ using fluorescence microscopy. Vacuoles were stained with FM4-64. Size bar, 2 μm, arrows show representative Ypt7 accumulations. (**D**) Quantification of Ypt7-positive dots in (**C**). Cells (n≥50) were counted from three independent experiments; (P-value ***<0.001 using two sample t-test). (**E**) Localization of mCherry-tagged Vps21 in cells expressing mNeon-Ypt7 and endogenously expressed wildtype or Mon1^Δ100^ using fluorescence microscopy. Size bar 2 μm, arrows show representative Vps21 accumulations. (**F**) Quantification of Vps21-positive dots in (**E**). Cells (n≥50) were counted from three independent experiments. (P-value ***<0.001 using two sample t-test). (**G**) Effect on cell growth by Mon1 truncations. Strains endogenously expressing wild-type or Mon1^Δ100^ were grown to the same OD_600_ in YPD media and spotted in serial dilutions onto agar plates containing YPD or YPD supplemented with 4 mM ZnCl_2_. Plates were incubated for several days at 30 °C. (**H**) A hydrophobic patch mutation in the N-terminal part of Mon1 affects GEF activity. Liposomes were loaded with 150 nM prenylated Ypt10 in the presence of 200 μM GTP and 1.5 mM EDTA. Nucleotide was stabilized using 3 mM MgCl_2_. 250 nM Mant-GDP loaded Ypt7:GDI were added and reaction was triggered by adding 12.5 nM wild-type (blue) or mutant complex expressing Mon1^L95A,L96A^ (green). Decrease in fluorescence detected over time is normalized to initial fluorescence. (**I**) Comparison of fold-change in GEF activity of wild-type and Mon1^L95A,L96A^ mutant GEF complexes on liposomes and in solution. For details of in solution GEF assay see method section. For the GEF assay on liposomes, k_obs_ of each curve was determined as described in the method section. K_obs_ values were normalized to the wild-type value. For in solution assays, k_cat_/K_m_ values were determined and normalized to the wild-type value. Bar graphs represent average foldchange and dots represent individual changes from three experiments. (P-value * p<0.05 using two sample t-test).

To test for conservation, we turned to the yeast Mon1-Ccz1 complex as in a genetically more amenable system. We deleted the first 100 residues in *S.c*. Mon1 (*S.c*. Mon1^Δ100^), which correspond to the region including the hydrophobic patch, (Fig. 1G, S2E) and analyzed the corresponding mutant for Ypt7 (yeast Rab7) and Vps21 (yeast Rab5) localization. In wildtype cells, endogenously expressed mNEON-Ypt7 localized to the vacuolar membrane and single dots. Cells expressing Mon1^Δ100^ had round vacuoles, suggesting sufficient Ypt7 activation. However, Ypt7 now localized more prominently to several perivacuolar dots in addition to its localization to the vacuolar membrane, suggesting an endosomal localization (Fig. 2C,D). We then analyzed the localization of the early endosomal Rab5-like Vps21 and observed that *S.c*. Mon1^Δ100^ cells had significantly less Vps21-positive dots than wildtype cells (Fig. 2E,F). This again suggests that the balance between early Rab5/Vps21-positive endosomes and late endosomal Rab7/Ypt7-positive structures is altered in Mon1^Δ100^ expressing cells. The overall expression of both Rab-GTPases was unaltered (Fig. S2F,G). To determine if the truncation of Mon1 affects cellular physiology, we grew cells in sequential dilutions on plates containing 4 mM ZnCl2 (Fig. 2G). *S.c*. Mon1^Δ100^ cells showed a slight growth defect in comparison to wild-type cells, indicating that *S.c*. Mon1^Δ100^ cells had problems with ion detoxification due to a defect in the endolysosomal pathway. To determine if *S.c*. Mon1^Δ100^ cells had also defects in autophagy, we followed transport of the cytosolic Pho8Δ60 reporter into the vacuole during nitrogen starvation (Duarte et al., 2022), but detected no difference in Pho8 activity of mutant versus wild-type cells (Fig. S2H).

We then asked if the N-terminal truncation of *S.c*. Mon1 also resulted in higher Mon1-Ccz1 activity as observed for the truncated *D.m*. Mon1. However, the N-terminal deletion in Mon1 did not yield sufficiently stable purified Mon1-Ccz1 complex. We therefore tested if a mutation in the corresponding hydrophobic motif present in the N-terminal domain (Fig. 1G; *S.c*. Mon1^L95A,L96A^) phenocopies the effect of the mutation in *D.m*. Mon1 (Fig. 1E). These mutations allowed purification of stable Mon1-Ccz1 (Fig. S2I), which had an increased Rab5-dependent activity on liposomes when compared to wildtype, but comparable GEF activity in solution (Fig. 2H,I, Table S3). However, this mutation in Mon1 did not impair growth of corresponding mutant strains on plates containing ZnCl2 (Fig. 3H), in agreement with the milder phenotype *in vitro* (Fig. 2H). Overall, these findings are consistent with the altered endocytic system of *Drosophila* nephrocytes that express the more active *D.m*. Mon1^Δ100^.

**Figure 3.**
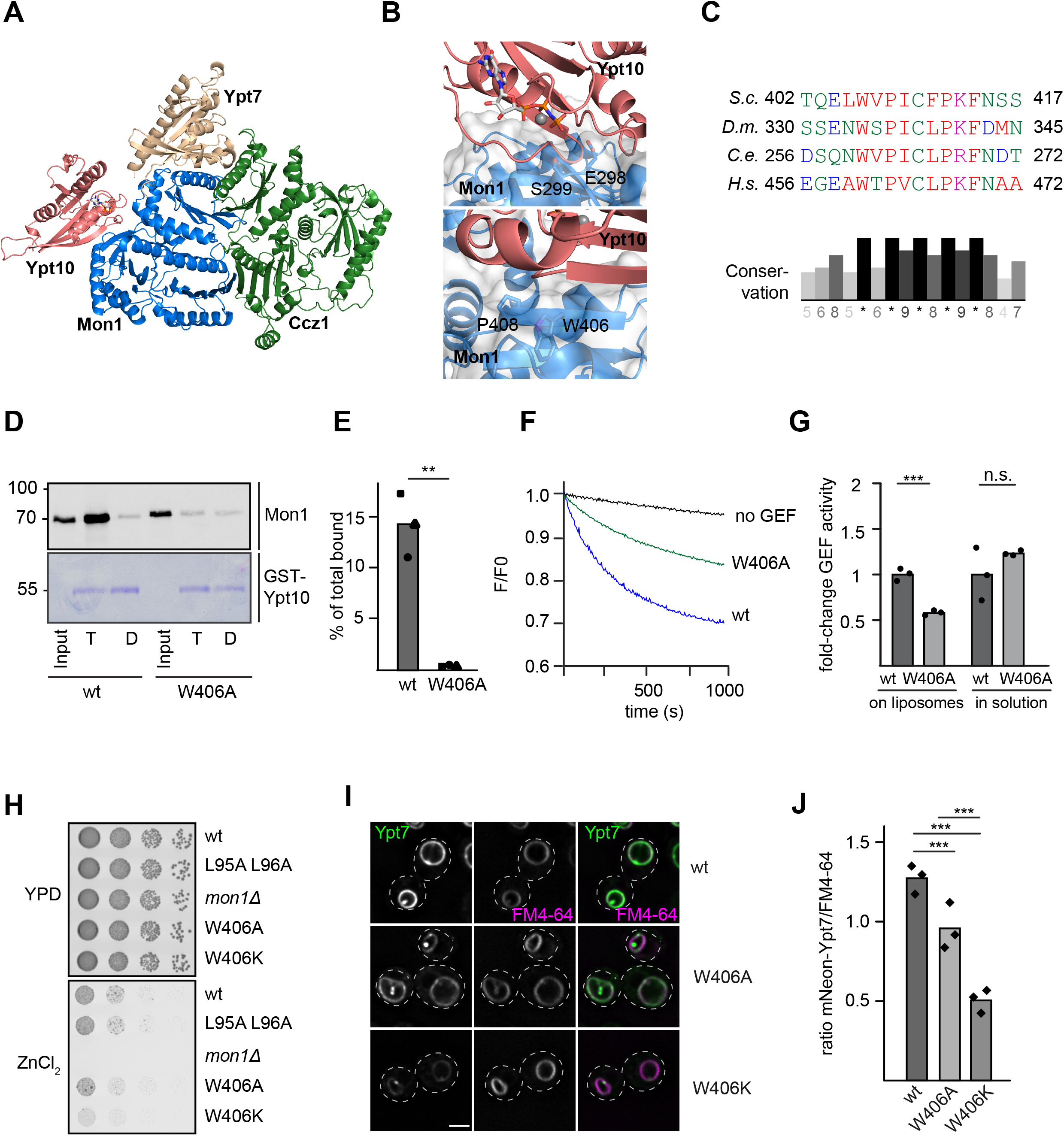
Identification of the Rab5 binding site in the Mon1-Ccz1 complex. (**A**) Identification of the putative Rab5 binding site in Mon1. Composite model of the *S.c*. Mon1 (blue)-Ccz1 (green)-Ypt7 (beige)-Ypt10 (pink) complex based on an Alphafold 2 prediction (Jumper et al., 2021; Evans et al., 2022) and the crystal structure of the catalytic core complex(PDB ID: 5LDD)(Varadi et al., 2021; Jumper et al., 2021; Kiontke et al., 2017). (**B**) Close-up view of the Mon1-Ypt10 binding interface. Colors are as in **A**. (**C**) Multiple sequence alignment of Mon1 β-strand in the modeled Rab5 binding region. Mon1 sequences from *Saccharomyces cerevisiae (S.c.)*, *Drosophila melanogaster (D.m.)*, *Caenorhabditis elegans (C.e.)* and *Homo sapiens (H.s.)* were aligned using Clustal omega web interface. Conservation was determined using Jalview. (**D**) Effect of the Mon1^W406A^ mutant on Rab5 binding. 150 μg purified GST-Ypt10 was loaded with GTP (T) or GDP (D) and incubated with 25 μg of either wild-type or Mon1^W406A^ GEF complex. Elution of bound GEF was performed with EDTA. 20 % of the eluate was analyzed together with 1 % input by Western blotting using an anti-Mon1 antibody. 2 % GST-Ypt10 were stained with Coomassie as loading control. (**E**) Quantification of bound GEF complex to Ypt10-GTP. Band intensity of Mon1 signal in elution fraction was measured using Fiji and compared to input. (P-value **<0.01 using two sample t-test). (**F**) Effect of the Mon1^W406^ mutation on Rab5-dependent GEF activity. 250 nM Mant-GDP loaded Ypt7:GDI were added and Rab activation was measured by fluorescence decrease over time. Liposomes were loaded with 150 nM prenylated recruiter Ypt10 using 200 μM GTP and 1.5 mM EDTA. Nucleotide was stabilized using 3 mM MgCl_2_. Reaction was triggered by adding 25 nM GEF complex. Decrease in fluorescence was normalized to fluorescence prior to GEF addition. (**G**) Comparison of fold-change in GEF activity of Mon1^W406A^ to Mon1^wt^ complex on liposomes and in solution. For details of in solution GEF assay see methods. For GEF assay on liposomes, k_obs_ of each curve was determined as described in the method section. K_obs_ values of mutant were normalized to the wild-type GEF complex value. For in solution assays, k_cat_/K_m_ values were normalized to the wild-type value. Bar graphs represent average fold-change and dots represent individual values from three experiments. (P-value ***<0.001, using two sample t-test). **(H)**Growth assay. Indicated Mon1 variants in a *mon1* deletion background were grown to the same OD_600_ in YPD media and then spotted in serial dilutions onto agar plates containing YPD or YPD supplemented with 4 mM ZnCl2. Plates were incubated for several days at 30 °C. **(I)**Localization of Ypt7 in wild-type and mutant strains. Plasmids encoding Mon1^wt^, Mon1^W406A^ or Mon1^W406K^ variants were expressed under their endogenous promoter in a *mon1* deletion strain. Vacuoles were stained with FM4-64. Images were deconvolved (SOftWoRx). Size bar, 2 μm. **(J)**Quantification of ratio between FM4-64 and mNeon-Ypt7 mean intensity signal on the vacuolar rim. Mean intensity signals were determined by a line profile across the vacuole. Cells (n≥50) were counted from three independent experiments. (P-value ***<0.01 using two sample t-test).

Taken together, these results suggest a regulatory role of the intrinsically disordered Mon1 N-terminal region including a short, conserved hydrophobic motif. This regulation is only observed in the Rab5-dependent GEF-assay on liposomes, suggesting that the N-terminal region either affects Rab5 binding to Mon1-Ccz1 or access of the substrate Ypt7 to the active site in a membrane context.

### Rab5 interacts with a conserved region of Mon1

To further elucidate the Rab5-dependent regulation of Mon1-Ccz1 GEF-activity, we searched for interaction sites between Rab5 and Mon1-Ccz1 using AlphaFold2. As we observed the strongest interaction between the Rab5-like Ypt10 and *S.c*. Mon1-Ccz1 in the past (Langemeyer et al., 2020), we used these proteins as templates for our modeling. Using AlphaFold2 multimer we generated a model for the *S.c*. Mon1-Ccz1 complex bound to Ypt7 (nucleotide-free) and Ypt10 (GTP-bound) (Fig.3A, S3A-C). In these models, Ypt10 was consistently positioned to interact with a beta-sheet in LD2 of Mon1 with high confidence (S3C). A similar binding mode is predicted for *D.m*. Rab5 to *D.m*. Mon1 (Fig. S3D,E) and observed for the interaction of Rsg1 to the Fuzzy subunit of CPLANE (Fig. S3F), further supporting the validity of the proposed interaction site. This region is next to the so-called elbow-loop of Mon1, which is involved in Mon1-Ccz1 GEF function and its interaction with Ypt7 (Kiontke et al., 2017). The loop residues E298 and S299 of Mon1 are predicted to interact with the P-loop and switch regions of Ypt10 (Fig. 3B, upper panel).

The proposed interface contains a highly conserved motif in LD2 of Mon1 with a hydrophobic residue (Fig. 3C) that corresponds to the surface-exposed W406 in *S.c*. Mon1 (Fig. 3B lower panel). Expression and purification of a corresponding Mon1^W406A^-Ccz1 complex resulted in a stable preparation (Fig. S3G). We then asked if the mutant complex was still able to interact with the Rab5-like Ypt10. We therefore bound GTP- or GDP loaded GST-Ypt10 on glutathione sepharose and added either purified wild-type or Mon1^W406A^ complex. Whereas wild-type Mon1-Ccz1 showed robust binding to Ypt10-GTP as shown before (Langemeyer et al., 2020), almost no Ypt10-binding was observed for the mutant complex, suggesting that the mutation strongly impaired Ypt10 binding (Fig.3D,E). In agreement, the purified Mon1^W406A^-Ccz1 complex had half the Rab5-dependent GEF-activity towards Ypt7 on membranes (Fig. 3F,G, Table S3). Importantly, GEF-activity was unperturbed in solution (Fig. 3G, Table S3). This suggests that Rab5 binding results in activation of Mon1-Ccz1 on membranes.

We wondered if deficient Rab5 binding of Mon1^W406A^ would affect Mon1-Ccz1 function *in vivo*.To address this, we transformed a *MON1* deletion strain with plasmids either carrying *MON1* or *MON1^W406A^*. Both alleles rescued the growth defect of the *MON1* deletion strain on medium containing ZnCl2 (Fig.3H). (Stepp et al., 1997). We then analyzed vacuolar morphology and localization of mNEON-tagged Ypt7, and observed an almost wild-type-like localization of Ypt7 and normal vacuole morphology in Mon1^W406A^ cells (Fig. 3I,J). However, mutation of W406 into a basic lysine residue in Mon1 resulted in far more dramatic phenotypes. Cells expressing Mon1^W406K^ had a strong decrease in vacuolar rim signal (Fig. 3I,J), consistent with a growth deficiency *in vivo* (Fig. 3H). The overall expression of Ypt7 was not affected in these strains (Fig. S3H) This suggests problems with correct spatiotemporal activation of Ypt7, possibly due to insufficient Mon1-Ccz1 activation by *S.c*. Rab5.

To test if the Rab5-binding site in Mon1 is conserved among species (Fig. 3B,C), we went back to the *Drosophila* system. The structural analysis of the *D.m*. Mon1-Ccz1-Bulli complex (manuscript in preparation) suggested a similar binding interface for *D.m*. Rab5 as identified for *S.c*. Mon1 (Fig.S3D,E). We therefore generated a corresponding Mon1^W334A^ mutation in the trimeric Mon1-Ccz1-Bulli complex and obtained a stable complex (Fig. 4A). Strikingly, this complex showed reduced activity in the Rab5-dependent GEF-assay (Fig. 4B,C, Table S3), but had normal GEF-activity in solution towards Rab7 (Fig. 4C, Table S3).

**Figure 4.**
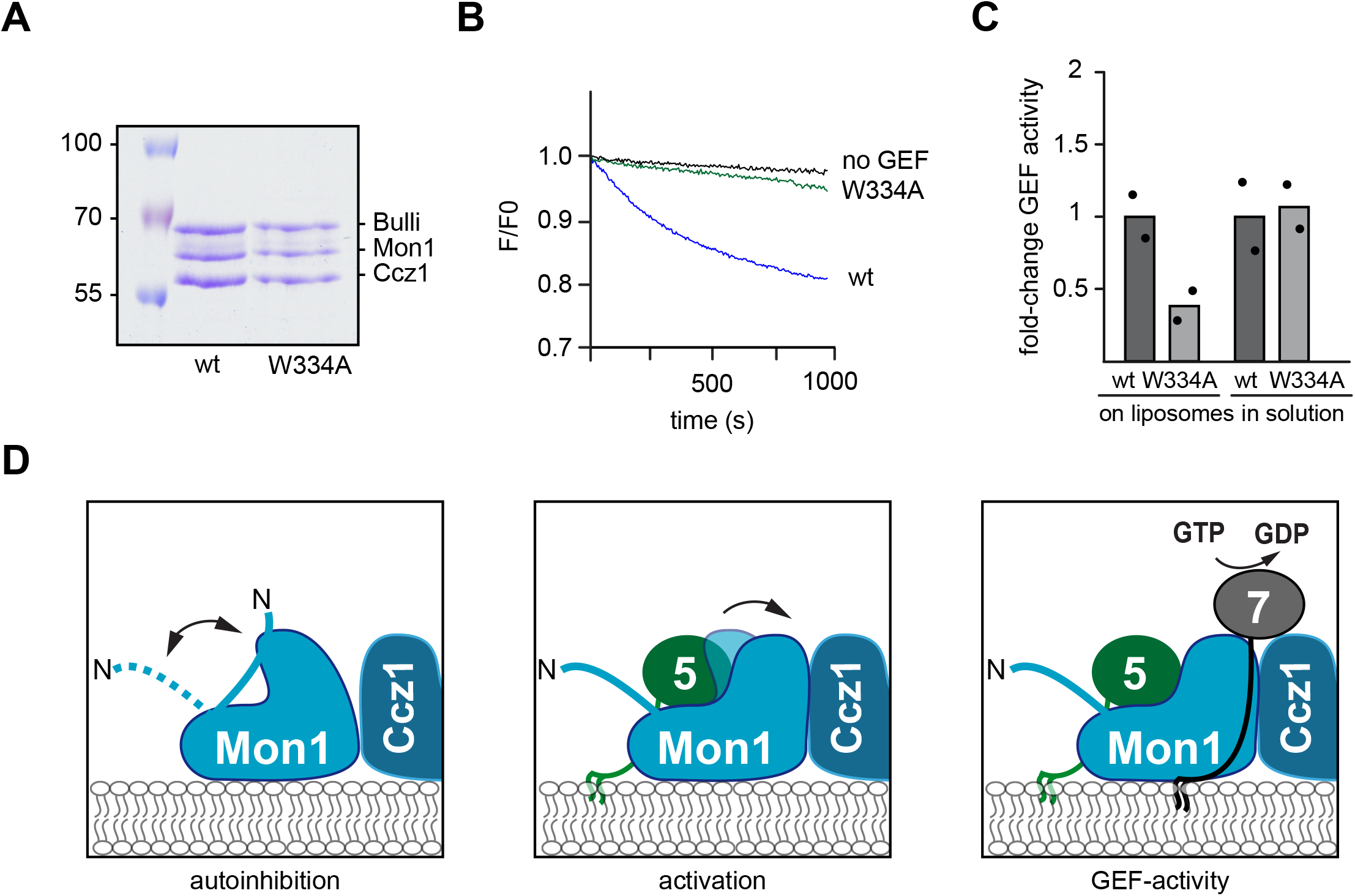
The Rab5 binding site in Mon1 is conserved. **(A)** Analysis of the *Drosophila* Mon1-Ccz1-Bulli complex with a Mon1^W334A^ mutation. Stability of wild-type and Mon1^W334A^ trimeric GEF complexes was analyzed by SDS-PAGE and Coomassie staining. **(B)**GEF activity of the wild-type and Mon1^W334A^ trimeric GEF complex. Rab activation was measured by fluorescence decrease over time. Liposomes were loaded with 150 nM prenylated Rab5 in the presence of 200 μM GTP and 1.5 mM EDTA, bound nucleotide was stabilized with 3 mM MgCl_2_. 250 nM Mant-GDP loaded Rab7:GDI complex were added and the reaction was triggered by adding 6.25 nM of the corresponding GEF complex. Decrease in fluorescence was normalized to fluorescence prior to GEF addition. **(C)**Fold-change in GEF activity of Mon1^W334A^ compared to wild-type complex on liposomes and in solution. For details see methods. Statistics were done as in Fig. 2I. Bar graphs represent average foldchange and dots represent individual values of two experiments normalized to corresponding wildtype. **(D)**Working model. The unstructured N-terminal part of Mon1 is folding back to the core of Mon1 resulting in autoinhibition of GEF-activity. Rab5 (green) binding to a conserved site activates Mon1-Ccz1 (blue) and drives nucleotide exchange of Ypt7. For details see text.

Taken together our results identify a conserved motif containing a hydrophobic tryptophan residue in Mon1 being involved in Rab5 binding and consequently in recruitment and activation of Mon1-Ccz1 (Fig. 3B, 4D).

## Discussion

Within this study, we focused on the regulation of the Mon1-Ccz1 complex. We here identify two conserved regions in Mon1, controlling the activity of the GEF complex on membranes. First, we show that deletion of the intrinsically disordered N-terminal part of Mon1 results in a two to three-fold more active GEF complex on membranes. This N-terminal part may thus function as an autoinhibitory loop for the complex. It is nevertheless required for Mon1-Ccz1 function *in vivo* as yeast cells as well as *Drosophila* nephrocytes lacking the Mon1 N-terminal have an altered Rab5 and Rab7 distribution along the endocytic pathway and physiological deficits. Second, we identify W406 as a key residue in Mon1 involved in Rab5 binding. Mutation of this residue results in poor binding to the Rab5-like Ypt10 in yeast, and strongly reduced Rab5-dependent GEF activity, both for the yeast and *Drosophila* Mon1-Ccz1 complex. Finally, we find a strong overlap of the identified site involved in Rab5 binding with the position of the non-canonical GTPase Rsg1 of the homologous CPLANE complex (Langousis et al., 2022), suggesting that the position of small regulatory GTPases to tri-longin domain (TLD)-GEF complexes may be conserved.

The Mon1-Ccz1 complex has a key role in the transition from early to late endosomes (Borchers et al., 2021). On endosomes, the GEF complex arrives at the same time as Rab7, which is accompanied by a sharp Rab5 to Rab7 transition (Rink et al., 2005; Poteryaev et al., 2010), and is later released from lysosomes (Yasuda et al., 2016). A similar release is observed from yeast vacuoles (Lawrence et al., 2014). Furthermore, Mon1-Ccz1 is required for Rab7 activation on autophagosomes (Gao et al., 2018; Hegedűs et al., 2016). Its targeting requires specific cues such as the recently identified amphipathic helix in Ccz1, which may recognize packaging defects within the membrane of autophagosomes (Herrmann et al., 2023). It is thus not unexpected that the complex is regulated at multiple sites. One of these is the N-terminal part of Mon1. Its removal results in a hyperactive complex and a defective endosomal system. This part of Mon1 is predicted to be intrinsically disordered, but also heavily phosphorylated (www.yeastgenome.org). Phosphorylation by one or multiple kinases or the accumulation of PI(3)P and Rab5 on endosomes may differentially control Mon1-Ccz1 activity (Fig. 4D). One possibility is that the Mon1 N-terminal controls Rab5 binding. On membranes without established Rab5 domains, the N-terminal region would then restrict Rab5 binding to the Mon1 subunit resulting in an autoinhibited GEF. Increasing local concentrations of Rab5 on growing endosomes, potentially accompanied by increasing PI(3)P concentrations produced by the Rab5 effector Vps34 kinase complex II (Tremel et al., 2021), would then act as maturation signal. Under these conditions, Rab5 binding and consequently Rab7 GEF activation could then take place. This activation would result in a sharp and timely regulated Rab5-to-Rab7 transition. The N-terminal region could also directly block the active site of the GEF complex, and Rab5 binding then in turn would allow structural rearrangements and release of the active site. However, we speculate that this region controls the access of Rab7 to Mon1-Ccz1, likely by restricting binding of the Rab7 HVD to the complex. According to our model, Rab7 would partition out of the GDI complex onto membranes, where it eventually encounters the GEF. A first interaction may require binding of the Rab7 HVD to the complex. If the N-terminal domain of Mon1 in an autoinhibited state would prevent binding of the HVD, Rab7 may not get efficient access to the active site.

Such a control of Rab access to the active site would be in part reminiscent of the gating mechanism of the TRAPP complexes (Thomas et al., 2019; Bagde and Fromme, 2022). These complexes share the same active site, but have distinct subunits in TRAPPIII and TRAPPII, which determine specificity. Here, the length of the HVD of Ypt1 targets it specifically to TRAPPIII, whereas the larger TRAPPII complex can only activate Ypt32 with its longer HVD (Thomas et al., 2019; Bagde and Fromme, 2022). It is likely that the interactions of the Rabs with the respective TRAPP complex is further regulated by interaction of the HVD with the core of the TRAPP complex. Future experiments will have to reveal if the Rab7-HVD is involved in interaction with Mon1-Ccz1 and how phosphorylation specifically controls Mon1-Ccz1 functions.

Multiple studies identified Rab cascades, where one Rab is required for the recruitment of the downstream GEF (Barr, 2013; Borchers et al., 2021; Hutagalung and Novick, 2011), yet mechanistic insight on how this works have been lacking. Here, we identify a region in Mon1 that can explain how Rab5 may not only recruit, but activate the Mon1-Ccz1 complex. Mutation of the conserved tryptophan residue within Mon1 strongly impairs binding to the Rab5-like Ypt10 and interferes with GEF activity of the yeast and *Drosophila* complex on membranes. The identified motif in Mon1 lies in a β-sheet of LD2 and in proximity to the elbow loop implicated in regulating the active site (Kiontke et al., 2017). This elbow loop was not resolved in the structure of the Mon1-Ccz1 complex in the absence of Ypt7 (Klink et al., 2022), suggesting that this region is involved in Ypt7 binding and activation. It is thus possible that Rab5 binding stabilizes the active site, and thus both recruits and activates the Mon1-Ccz1 complex as revealed in our *in vitro* assays (Langemeyer et al., 2020; Herrmann et al., 2023). Curiously, the non-canonical small GTPase Rsg1 can bind to the tri-longin CPLANE complex, and here occupies the same site on the Fuzzy subunit, where Rab5 binds Mon1 (Langousis et al., 2022). We thus speculate that Rsg1 may also activate the CPLANE complex.

Our data reveal that the targeting and activation mechanism of the Rab7 GEF is conserved. However, we still understand very little of the precise regulation. Our recent analysis of the lipid and protein requirements of the Mon1-Ccz1 complex showed that targeting to autophagosomes requires an amphipathic helix in Ccz1 and Atg8, whereas targeting to endosomes relies on negative charges and Rab5. Regulation may occur by distinct posttranslational modifications, likely phosphorylation by kinases such as the casein kinase Yck3 (Lawrence et al., 2014; Langemeyer et al., 2020). Other kinases may affect Mon1-Ccz1 in autophagy. One obvious regulatory site is the N-terminal segment of Mon1. Here, phosphorylation and dephosphorylation may occur by multiple kinases and phosphatases, depending on the organelleMon1-Ccz1 complex is targeted to. As organelle maturation depends on the spatiotemporal activation of the GEFs, it is crucial to understand the regions involved in targeting and regulation. Our analysis of Mon1-Ccz1 may thus help to understand the underlying mechanisms guiding Rab exchange and organelle maturation.

## Material & Methods

### Strains and Plasmids

Strains used in this study are listed in Table S1. Genetic manipulation of yeast strains including deletion and tagging of genes was performed using a PCR-product- and homologous recombination-based approach described in (Janke et al., 2004). Plasmids used in this study are listed in Table S2.

### Expression and purification of Rab GTPases and prenylation machinery components

GST-TEV-Rab5, GST-TEV-Rab7, GST-TEV-Ypt10, GST-TEV-Ypt7 and the components of the prenylation machinery, Bet4, His-TEV-Bet2, His-Sumo-*D.m*. GDI, GST-PreSc-Gdi1 and Mrs6-His were expressed and purified as before (Table S2)(Langemeyer et al., 2020). *E. coli* BL21 Rosetta cells were transformed with 100 ng plasmid DNA and grown in LB media to an OD_600_=0.6 at 37 °C. Protein expression was induced using 0.25 mM IPTG for 14 h at 16 °C. Cells were harvested and lysed in 50 mM HEPES-NaOH, pH 7.5, 150 mM NaCl, 1.5 mM MgCl_2_, 1 mM DTT, 1 mM PMSF and 0.05x Protease inhibitor cocktail (PIC; a 20X stock solution contained 2 μg/ml Leupeptin, 10 mM 1,10-Phenanthroline, 10 μg/ml Pepstatin A and 2 mM Pefablock) using the microfluidizer (Microfluidics, Westwood, USA). Yeast and *Drosophila* Rab GTPases were either purified as GST-fusion constructs by Glutathione elution for usage in GST pull downs or as Tag-free version by overnight cleavage with TEV protease. *S.c*. GST-PreSc-Gdi1 was purified by PreScission protease (PreSc) cleavage. Bet4 His-TEV-Bet2, Mrs6-His, and His-Sumo-*D.m*. GDI were added to Ni-NTA agarose from cleared lysates and eluted with 300 mM imidazole in elution buffer. The GST-PreSc-Gdi1 construct was cleaved for 2 h at 16 °C in the presence of 0.5 mM DTT. Proteins eluted with imidazole or glutathione were dialyzed overnight against buffer with 50 mM HEPES-NaOH, pH 7.5, 150 mM NaCl and 1.5 mM MgCl_2_ with one buffer exchange. To remove the His-Tag from His-Sumo-*D.m*. GDI, the dialyzed protein was incubated with SUMO protease for 2 h at 4 °C, and protease was then removed using Ni-NTA beads. Proteins were aliquoted, snap frozen in liquid nitrogen, and stored at −80 °C. The purity and efficiency of the purifications was analyzed by SDS gel electrophoresis.

### Expression and purification of *Drosophila* GEF complexes in Sf21 cells

*Drosophila* Mon1-Ccz1 and Mon1-Ccz1-Bulli were expressed using the biGBac system (Weissmann et al., 2016) and purified as described (Langemeyer et al., 2020). Briefly, Sf21 cells were grown as a monolayer culture in Insect-XPRESS Protein-free Insect Cell Medium (Lonza) in standard T175 culture flasks at 27 °C. Viral infection was performed for 72 h. Cells were then harvested at 500 *g* for 5 min, resuspended in buffer containing 50 mM HEPES-NaOH, pH 7.5, 300 mM NaCl, 1 mM MgCl_2_, 10% (v/v) glycerol, 1 mM PMSF, and 0.05x PIC and lysed using the microfluidizer (Microfluidics,Westwood, USA) or via homogenization. The cleared lysate was incubated for 2 h with Glutathione 4B Sepharose (GE Healthcare, Germany) followed by washing of the Sepharose with 50 mM HEPES-NaOH, pH 7.5, 300 mM NaCl, 1,5 mM MgCl_2_, and 10% glycerol and overnight cleavage of protein in the presence of 1 m DTT and 0.4 mg/ml PreScission protease. The next morning, protein was eluted and concentrated to 500 μl using a Vivaspin6 10,000 MVCO centrifugal concentrator (Sartorius, Germany). The concentrated sample was subjected to size exclusion chromatography using an Äkta FPLC UPC-900 liquid chromatography system (GE Healthcare, Solingen, Germany) equipped with a Superdex 200 increase 10/300 GL column (GE Healthcare, Solingen, Germany) and buffer containing 50 mM HEPES-NaOH, pH 7.5, 300 mM NaCl, 1 mM MgCl_2_, and 10 % (v/v) glycerol. Peak fractions were collected and analyzed by SDS-PAGE.

### Tandem-Affinity Purification

Purification of yeast Mon1-Ccz1 was essentially performed as described (Nordmann et al., 2010) with all centrifugation steps conducted at 4 °C. Three liters of culture were grown in YPG to an OD_600_=2-3 at 30 °C. Cells were harvested at 4000 *g*, and once washed in ice cold H2O. Pellets were resuspended in equal amounts of buffer containing 50 mM HEPES-NaOH, pH 7.4, 150 mM NaCl, 1.5 mM MgCl_2_, 5 % (v/v) glycerol, 1x FY, 0.5 mM PMSF and 0.5 mM DTT and were dropwise snap frozen in liquid nitrogen. Drops were then subjected to cryomill lysis, and powder was resuspended in equal amount of buffer on a nutator. Lysate was centrifuged for 10 min at 3200 *g* and then for 70 min at 125,000 *g*. Supernatant was incubated for 1.5 h at 4 °C with IgG Sepharose (Cytiva) equilibrated with buffer containing 50 mM HEPES-NaOH, pH 7.4, 150 mM NaCl, 1.5 mM MgCl_2_ and 5 % glycerol. Beads were washed extensively, and protein was cleaved over night at 4 °C in buffer with 1 mM DTT and TEV protease. Proteins containing fractions were analyzed on SDS gels followed by Coomassie staining.

### *In vitro* prenylation of yeast and *Drosophila* Rab GTPases

Prenylated yeast and *Drosophila* Rab-REP and Rab-GDI complexes were generated as described in (Langemeyer et al. 2018; Langemeyer et al. 2020; Thomas and Fromme 2016). For the generation of *Drosophila* Rab7-GDI, the yeast prenylation machinery and the *Drosophila* GDI were used. Rab proteins were either loaded with GDP (REP-complexes, Sigma Aldrich, Germany) or MANT-GDP (GDI-complexes, Jena Bioscience, Germany) prior to the prenylation reaction in prenylation buffer containing 50 mM HEPES-NaOH, pH 7.4, 150 mM NaCl, and 1.5 mM MgCl_2_.

### In solution and membrane-associated fluorescent nucleotide exchange assays

In solution nucleotide exchange factor assays were essentially performed as in (Kiontke et al. 2017) with minor changes. Purified Ypt7/Rab7 were loaded with MANT-GDP (GDI-complexes, Jena Bioscience, Germany) in the presence of 20 mM HEPES/NaOH, pH 7.4, and 20 mM EDTA for 30 min at 30 °C. Bound nucleotide was stabilized with 25 mM MgCl_2_. 2 μM Rab were incubated with varying amounts of GEF in a SpectraMax M3 Multi-Mode Microplate Reader (Molecular Devices, San Jose, USA) and after base line stabilization, nucleotide exchange was triggered by 0.1 mM GTP (for *D.m*.GEFs) or 1 mM GTP (*S.c*. GEFs). MANT-GDP release following Rab activation was monitored over time using an excitation wavelength of 355 nm and emission at 448 nm. Data after 10 min initial signal decrease were fitted against a first-order exponential decay using OriginPro9 software (OriginLab Corporation, Northampton, USA) and k_obs_ (s^-1^) was determined as 1/t1. k_obs_ was then plotted against the GEF concentration, and k_cat_/K_M_ (M^-1^s^-1^) was derived as the slope of the resulting linear fit.

The liposome-based GEF assays were performed as described (Langemeyer et al. 2020) with liposomes with a vacuolar mimicking composition (Zick and Wickner, 2014) with 47.6 mol % dioleoyl phosphatidylcholine (DLPC 18:2 18:2), 18 % dioleoyl phosphatidylethanolamine (DLPE 18:2 18:2), 18 % soy phosphatidylinositol (PI), 1 % diacylglycerol (DAG 16:0 16:0), 8 % ergosterol, 2 % dioleoyl phosphatidic acid (DLPA 18:2 18:2), 4.4 % dioleoyl phosphatidylserine (DLPS 18:2 18:2), 1 % dipalmitoyl PI(3)phosphate (PI(3)P diC16) (Life Technologies) and extruded to 400 nm using a polycarbonate filter and a hand extruder (Avanti Polar Lipids, Inc). Liposomes were decorated with indicated amounts of prenylated recruiter GTPase in the presence of 200 μM GTP and 1.5 mM EDTA for 15 min at RT. The loading reaction was stopped with 3 mM MgCl_2_, and the mix was transferred to a half micro cuvette 109.004F-QS with 10 x 4 mm thickness (Hellma). 250 nM Rab7/Ypt7-GDI were added and the cuvette was filled up to 800 μl with prenylation buffer omitting the volume of the GEF. The cuvette was placed in a fluorimeter (Jasco, Gross-Umstadt, Germany) at 30 °C with the wavelengths mentioned above. After baseline stabilization, the indicated concentration of GEF was added to trigger nucleotide exchange, and fluorescence decrease was monitored over time. Data were fitted against a first-order exponential decay using OriginPro9 software (OriginLab Corporation, Northampton, USA) and k_obs_ (s^-1^) was determined as 1/t1. All kinetic parameters of used GEF complexes are listed in Table S3.

### GST Rab pull downs

75 μg GST-tagged Rab GTPase was loaded with either 10 mM GDP or GTP (Sigma Aldrich, Germany) in the presence of 20 mM EDTA and 50 mM HEPES-NaOH, pH 7.4 at 30 °C for 30 min. The nucleotide was stabilized using 25 mM MgCl_2_, and protein was incubated for 1 h at 4 °C with 30 μl GSH sepharose (Cytiva, Germany) equilibrated with 50 mM HEPES-NaOH, pH 7.4, 150 mM NaCl, 1.5 mM MgCl_2_ in the presence of 7 mg/ml BSA. Then the beads were spun for 1 min, and the supernatant was discarded. Next, 25 μg Mon1-Ccz1, 7 mg/ml BSA and 1 mM nucleotide were added, and the tube was filled up to 300 μl with pull down buffer containing 50 mM HEPES-NaOH, pH 7.4, 150 mM NaCl, 1 mM MgCl_2_, 5% (v/v) Glycerol and 0.1% (v/v) Triton X-100, and incubated 1.5 h at 4 °C on a turning wheel. Then beads were washed 3x using pull down buffer. Subsequently, bound protein was eluted from the beads for 20 min at RT in a turning wheel using 300 μl elution buffer with 50 mM HEPES-NaOH, pH 7.4, 150 mM NaCl, 20 mM EDTA, 5% (v/v) Glycerol and 0.1% (v/v) Triton X-100. Eluted fraction was TCA-precipitated, and 40 % of it was analyzed by SDS page and Western Blot together with 1 % input sample. Mon1 was detected using a Mon1-antibody (Ungermann lab) and a fluorescence-coupled secondary antibody ((#SA5-35571 Thermo Fisher Scientific). For the Rab GTPase as loading control, 1x Laemmli buffer was added to the GSH beads after elution, and samples were boiled for 10 min at 95 °C. 0.4 % of each loading control was loaded onto SDS page and stained with Coomassie Brilliant blue G250.

### Pho8Δ60 assay

The assay was essentially performed as in (Guimaraes et al. 2015). Yeast strains expressing a genetically truncated version of the *PHO8* gene (Pho8Δ60) were grown in YPAD media overnight at 30 °C. The next morning, cells were diluted to an OD_600_=0.2 in 10 ml and grown until logarithmic phase. Then cells were centrifuged, washed in starvation media (0.17% yeast nitrogen base without amino acids or ammonium sulfate and 2 % glucose), and starvation was induced for indicated time periods. Then 5 OD equivalents of cells were harvested and washed in H2O. As control, 5 OD equivalents of cells grown in YPAD were treated the same. Pellets were resuspended in buffer 20 mM PIPES, pH 6.8, 0.5% Triton X-100, 50 mM KCl, 100 mM potassium acetate, 10 mM MgCl_2_, 10 μM ZnSO_4_ and 2 mM PMSF and cells were lysed by glass bead lysis in the FastPrep-24 (MP Biomedicals). Pho8 activity was monitored by incubation of 200 μl lysate with 50 μl buffer 250 mM Tris–HCl, pH 8.5, 0.4% Triton X-100, 10 mM MgCl_2_, 10 μM ZnSO_4_ and 1.25 mM p-nitrophenyl phosphate and the colorimetric reaction was measured in a SpectraMax M3 Multi-Mode Microplate Reader (Molecular Devices, San Jose, USA) at OD400.The enzymatic activity was calculated by the linear method. The slopes of the experimental samples were normalized to the ALP activity of WT cells under nutrient rich conditions and displayed in arbitrary units (A.U.).

### Analysis of yeast and *Drosophila* protein expression

To test the expression of Rab GTPases Vps21 and Ypt7 in yeast cells, 4 OD units were harvested and resuspended in buffer containing 0.2 M NaOH and 30 mM β-mercaptoethanol. Proteins were precipitated by adding TCA to a final concentration of 15 % and analyzed via SDS PAGE and Western Blotting. Antibodies against Vps21 (1:1000) and Ypt7 (1:3000) were generated in the Ungermann lab, the Tom40 antibody was a gift of the Neupert lab. Primary antibodies were detected with secondary antibody goat anti-Rabbit IgG (H+L) Secondary Antibody, DyLight™ 800 4X PEG (#SA5-35571, Thermo Fisher Scientific, 1:10000) or DyLight™ 680 (#35568, Thermo Fisher Scientific,1:10000). The expression of UAS-driven wild-type Mon1::HA and truncated Mon1^Δ100^::HA in transgenic flies using the ubiquitous *daughterless*-GAL4 driver was verified by western blot of *Drosophila* whole cell lysate from two female flies. Actin staining was used as a loading control. As a primary antibody rabbit α-HA (1:1000, Sigma-Aldrich, H6908) and mouse α-Actin (1:10, DSHB, JLA20) was used, combined with a secondary antibody α-rabbit-alkaline phosphatase (1:10,000, Sigma-Aldrich, A3687) and α-mouse-alkaline phosphatase (1:10,000, Sigma-Aldrich, A3562).

### Fluorescence microscopy

For fluorescence microscopy, cells were grown overnight in synthetic media containing 2 % (w/v) glucose and essential amino acids (SDC+all). In the morning, cells were diluted to an OD_600_=0.1 and grown to logarithmic phase. Vacuoles were stained with FM4-64 (Thermo Fisher Scientific) where indicated. 1 OD equivalent of cells was resuspended in 50 μl SDC+all containing 30 μM FM4-64 and incubated at 30 °C for 20 min. Cells were washed twice with fresh media and incubated with 500 μl fresh SDC+all for another 45 minutes at 30°C. Cells were imaged on an Olympus IX-71 inverted microscope (DeltaVision Elite) equipped with a 100x NA 1.49 objective, a sCMOS camera (PCO, Kelheim, Germany), an InsightSSI illumination system, SoftWoRx software (Applied Precision, Issaquah, WA) and GFP, mCherry and Cy5 filters. Stacks with 0.35 μm spacing were taken. Where indicated, deconvolution was performed. Microscopy images were processed as single slices and quantified using Fiji software (National Institutes of Health, Bethesda, MD).

### Growth test

Yeast cells were incubated in YPD media overnight at 30 °C. In the morning, cultures were diluted and grown to logarithmic phase at 30 °C. Then cells were diluted to an initial OD_600_=0.25 in YPD, and spotted in 1:10 serial dilutions onto control and selection plates and incubated for several days at the indicated temperature. Each day, plates were imaged to monitor growth.

### Fly stocks

The following fly stocks were obtained from the Drosophila stock center at Bloomington: *da*-GAL4 (RRID:BDSC_55850) and *w*^1118^ (RRID:BDSC_5905). *handC*-GAL4 was previously generated by us (Sellin et al., 2006). The UAS-Mon1::HA line was obtained from T. Klein, Düsseldorf, Germany (Yousefian et al., 2013) and the UAS-Mon1^Δ100^::HA line was generated in this study. Fly husbandry was carried out as described previously (Wang et al., 2012).

### Generation of the transgenic UAS-Mon1^Δ100^::HA line

To generate a Mon1^Δ100^::HA construct in a *Drosophila* vector suitable for establishing transgenic fly lines, a Mon1^Δ100^-cDNA was used as a PCR template (Source DNA clone IP03303 from the *Drosophila* Genomics Resource Center (DGRC, Indiana, USA)). Via overlap PCR, the Mon1^Δ100^ construct was fused to an HA-tag following by a PCR to generate restriction sites (XhoI and EcoRI) for insertion into an injection plasmid. Therefore, the following PCR primers were used: 5’ TACCGGAATTCATGGAGGAGGAATACGATTACCAGC ‘3 (forward) and 5’ TACCGCTCGA GTTAAGCGTAATCTGGAACG ‘3 (reverse). The PCR amplicon was purified, digested with EcoRI and XhoI and cloned into pYED (Paululat and Heinisch, 2012). The UAS-Mon1^Δ100^::HA expression plasmid was injected into RRID:BDSC_24749 for integration at 86Fb on the 3. chromosome (Bischof et al., 2007). A commercial service was used for establishing transgenic fly lines (BestGene, Chino Hills, CA, USA).

### FITC-Albumin Uptake

FITC-albumin uptake assays were performed in pericardial nephrocytes as described previously with minor modifications (Dehnen et al., 2020). Briefly, 3^rd^ instar larvae, in which Mon1::HA or Mon1^Δ100^::HA was expressed in nephrocytes using *handC-GAL4* as a driver, were fixed ventral side upwards on Sylgard 184 silicone elastomer plates using minute pins. Specimens were always covered by artificial hemolymph. All internal organs except for the pericardial cells and associated tissue (e.g. heart, alary muscles) were removed. Preparation buffer was replaced by fresh artificial hemolymph buffer containing 0.2 mg/ml FITC-albumin (A9771, Albumin–fluorescein isothiocyanate conjugate, MW: 66 kDa; Sigma Aldrich, Munich, Germany), and specimens were incubated in the dark for 5 min or 10 min. Uptake was stopped by fixation in 8 % paraformaldehyde (PFA) in phosphate-buffered saline (PBS) followed by two washing steps with PBS. Subsequently, tissues were embedded in Fluoromount-G mounting medium containing DAPI (Thermo Fisher, Waltham, USA) for microscopic analysis. Uptake efficiency was quantified by imaging respective nephrocytes (LSM800, Zeiss, Jena, Germany). The mean pixel-intensity measurement function provided by the Fiji software package (National Institutes of Health, Bethesda, MD) was used to quantify uptake efficiency (Schindelin et al., 2012). Mean pixel intensity was calculated in relation to the perimeter of the cell.

### Immunohistochemistry

3^rd^ instar larvae expressing *handC* driven wild-type Mon1::HA and truncated Mon1^Δ100^::HA were dissected in PBS and fixed with 4 % paraformaldehyde (PFA) in PBS for one hour at RT. After three washing steps of 10 min, specimens were permeabilized with 1 % Triton X-100 in PBS for one hour at RT, followed by three further washing steps with BBT (0.1 % BSA and 0.1 % Tween-20 in PBS) for 10 min each. Subsequently, specimens were incubated for 30 min in a blocking solution containing 1 % BSA and 0.1 % Tween-20 in PBS followed by incubation with the primary antibodies (rabbit anti-Rab5, 1:250, Abcam 3126, Cambridge, UK; mouse anti-Rab7, 1:10, Developmental Studies Hybridoma Bank, University of Iowa, USA) in BBT overnight at 8 °C. Samples were rinsed three times with BBT (10 min, RT), blocked with blocking solution for 30 min and incubated with secondary antibodies (antirabbit Alexa Fluor 488, 1:200, Jackson ImmunoResearch Laboratories, Inc (Code Number: 111-545-006); anti-mouse Cy3, 1:200, Dianova GmbH) in BBT for two hours at RT followed by three washing steps with BBT. Samples were embedded in Fluoromount-G mounting medium containing DAPI (Thermo Fisher, Waltham, USA). Confocal images were captured with a laser scanning microscope (LSM800, Zeiss, Jena, Germany). Image processing was done with Fiji and Affinity Photo (Serif). The Pearson Correlation Coefficient was calculated using the ZEN blue edition software.

### Transmission electron microscopy

Briefly, specimens were prepared in PBS and subsequently fixed for 4 h at RT in fixative (2% glutaraldehyde (Sigma, Germany)/ 4% paraformaldehyde (Merck, Germany) in 0.05 M cacodylate buffer pH 7.4). Next, specimens were post-fixed for 2 h at RT in 1% osmium tetroxide in 0.05 M cacodylate buffer pH 7.4 (Merck, Germany) and dehydrated stepwise in a graded ethanol series followed by 100% acetone. Subsequently, specimens were embedded in Epon 812 and polymerised for 48 h at 60 °C. Ultrathin sections (70 nm) were cut on an ultramicrotome (UC6 and UC7 Leica, Wetzlar, Germany) and mounted on formvar-coated copper slot grids. Sections were stained for 30 minutes in 2% uranyl acetate (Sciences Services, Germany) and 20 minutes in 3 % lead citrate (Roth, Germany). A detailed protocol for processing nephrocytes for TEM analysis can be found elsewhere (Psathaki et al., 2018). All samples were analyzed at 80 kV with a Zeiss 902, and Zeiss LEO912 and at 200 kV with a Jeol JEM2100-Plus transmission electron microscope (Zeiss, Jena, Germany; Jeol, Tokyo, Japan).

### Modeling of the yeast Mon-Ccz1 complex bound to Ypt7 and Ypt10

With the sequences of Mon1, Ccz1, Ypt7 and Ypt10, an initial model of the tetrameric complex was generated using AlphaFold2 multimer (Evans et al., 2022). The switch regions of nucleotide-free Ypt7 were manually edited based on the crystal structure of the catalytic MC1-Ypt7 core complex (Kiontke et al., 2017) and GTP and Mg^2+^ were added to the Ypt10 nucleotide binding pocket. Only regions that were modeled with high confidence are shown in the figures.

## Acknowledgements

We thank Kerstin Etzold, Angela Perz and Kathrin Auffarth for expert technical assistance. This work was supported by a grant from the Deutsche Forschungsgemeinschaft (DFG) to Christian Ungermann and Achim Paululat (UN111/9-2; PA517/12-2), and by the SFB 944 and SFB 1557 (to C. Ungermann, A.Paululat and D. Kümmel). We also acknowledge the support of the Open Access Publishing Fund of Osnabrück University.

