## Supplementary figures and images for "Regulatory sites in the Mon1-Ccz1 complex control Rab5 to Rab7 transition and endosome maturation"

### Figure S1

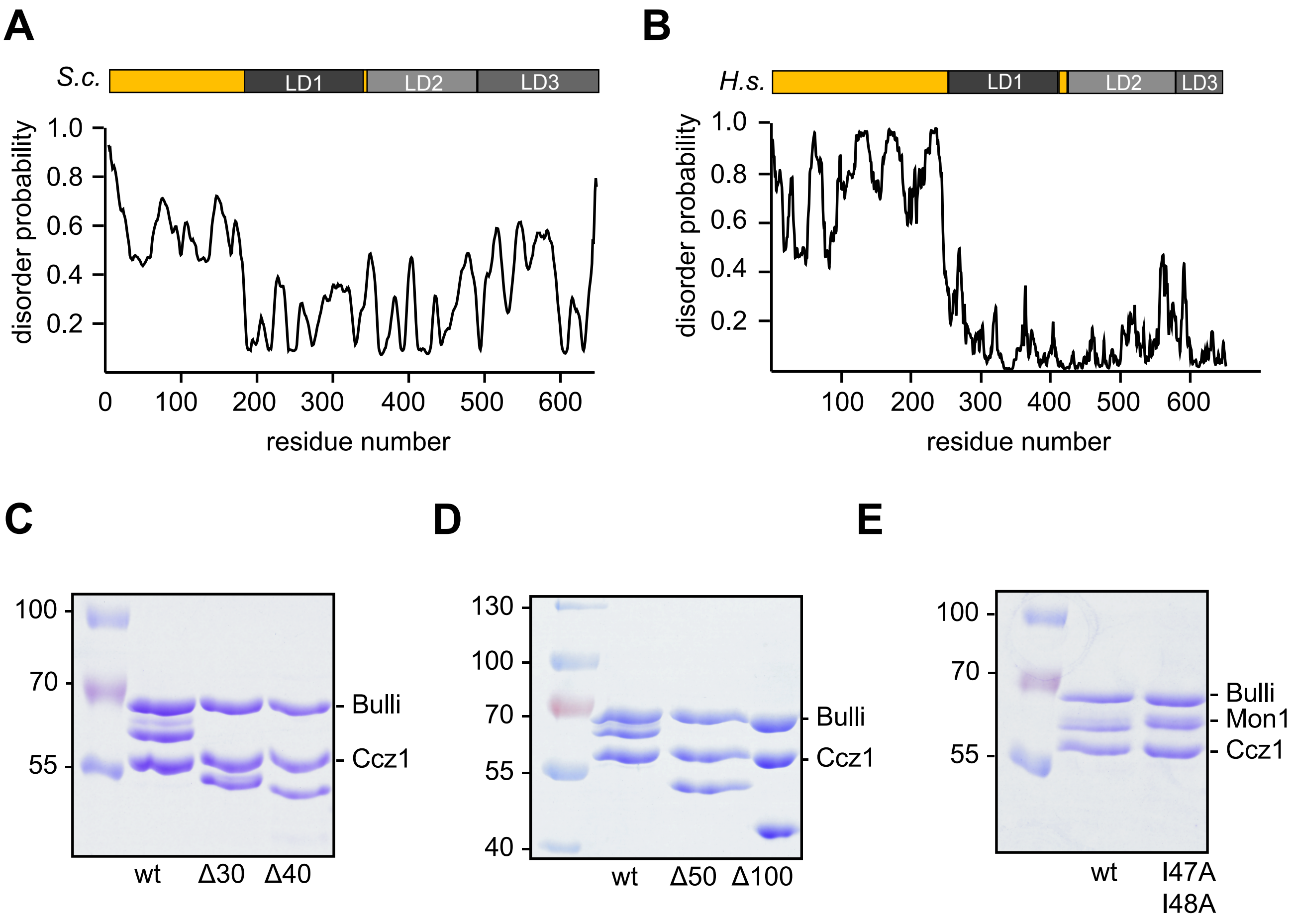

### Figure S2

Figure S2, Borchers et al.

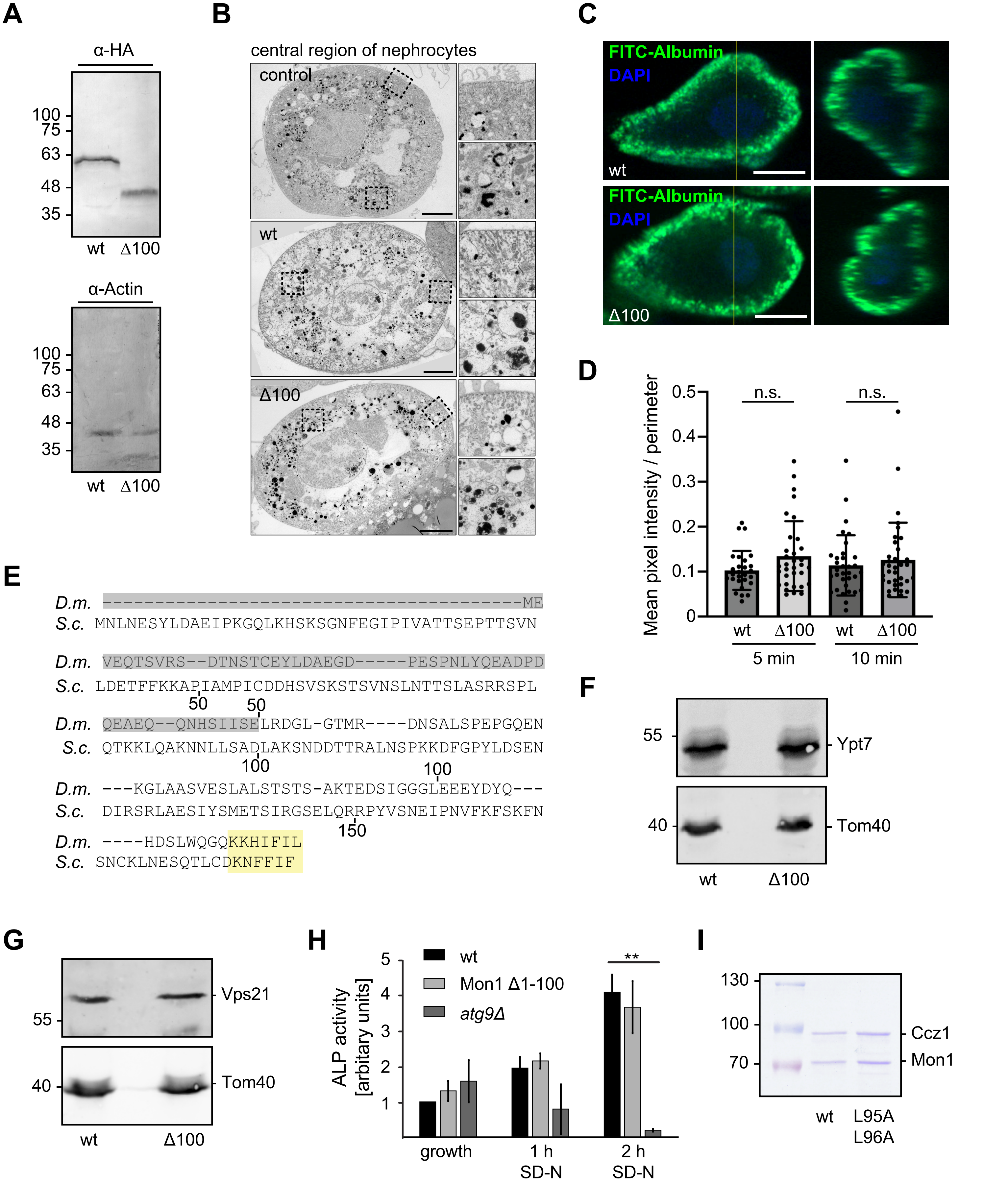

### Figure S3

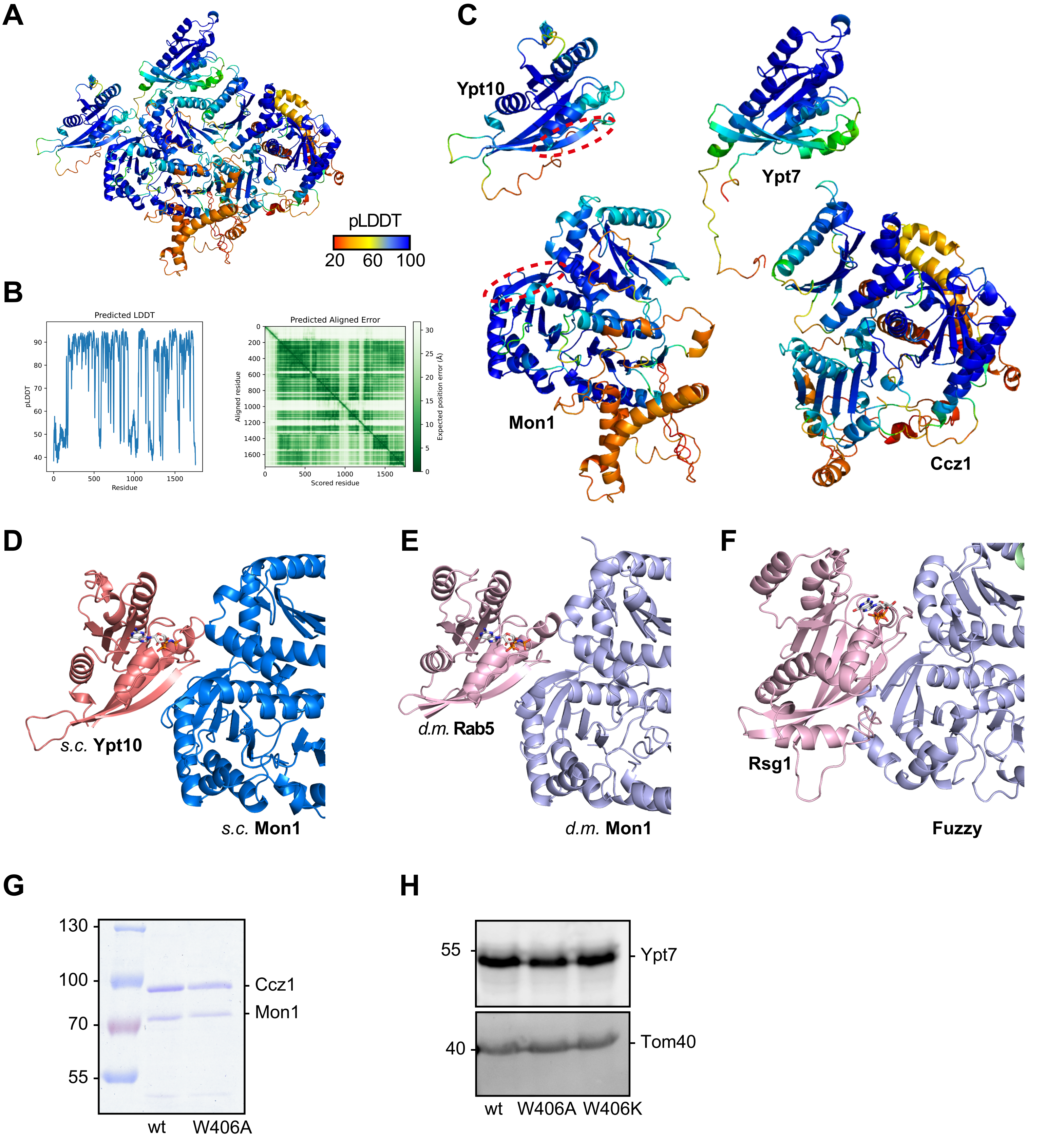
