## Supplemental Figure Legends for "Regulatory sites in the Mon1-Ccz1 complex control Rab5 to Rab7 transition and endosome maturation"

**Figure S1. Analysis of the Mon1 N-terminal region. (A,B)** The N-terminal regions of *S.c.* and *H.s.* Mon1 are disordered. Disorder probability of each residue of *S.c.,* (A) and *H.s*., (B) Mon1 was determined using IUPred2A web interface (Erdős and Dosztányi, 2020; Mészáros et al., 2018). Longin domains (LD) 1-3 are highlighted in grey shades. Values >0.5 are considered as disordered. **(C-E)** Expression of truncated Mon1 (Mon1^Δ1-30^, Mon1^Δ1-40^, **C**), and (Mon1^Δ1‑50^, Mon1^Δ1-100^, **D**) and of hydrophobic patch mutant (Mon1^I47, 48A^, **E**) does not affect complex stability of Trimeric Mon1-Ccz1-Bulli complex. GEF complexes were purified as described in the method section and analyzed by SDS-PAGE and Coomassie staining.

**Figure S2. Analysis of N-terminal mutations in Mon1.** (**A**) Protein levels of wild-type Mon1 and Mon1^∆100^ in adult female flies using the *daughterless-*GAL4 driver. Mon1 was detected via that HA-tag using α-HA (rabbit). Actin staining with α‑Actin (mouse) served as loading control. (**B**) TEM images of control nephrocytes (3^rd^ instar larvae from the crossing *handC*-GAL4 crossed to *white*^1118^) and nephrocytes from 3^rd^ instar larvae expressing wildtype or Mon1^∆100^ under control of the *handC*-GAL4 driver. Enlargements show the labyrinth channel system with slit diaphragm and clathrin coated vesicles. Further, endocytic vesicles with electron dense material are shown. (**C**) Uptake of FITC-Albumin (molecular weight of approximately 66 kDa) within nephrocytes from 3^rd^ instar larvae expressing wild-type Mon1 or Mon1^∆100^ under control of the *handC*-GAL4 driver. DAPI was used to visualize nuclei. Right panel shows an orthogonal view of the cells at the position marked with a yellow line. Size bar, 10 µm. (**D**) Quantification of FITC-Albumin uptake in nephrocytes for 5 min and 10 min. Regions of interest were analyzed for the mean pixel intensity in relation to the perimeter of the cell. For wild-type Mon1, 28 cells from 10 animals (5 min) and 33 cells from 11 animals (10 min) were quantified. For Mon1^∆100^, 33 cells from 11 animals (5 min) and 35 cells from 12 animals were quantified (10 min). (P‑value n.s.>using two sample t-test). **(E)** Alignment of *D.m.* and *S.c.* Mon1 N-terminal region. *D.m.* Mon1^Δ50^ (grey) corresponds to truncation of residues 1-100 in the *S.c.* protein. The beginning of LD1 is marked in yellow. Alignment was done with Clustal omega web interface (Sievers et al., 2011; Goujon et al., 2010). **(F,G)** Expression control of endosomal Rab GTPases Ypt7 (F) and Vps21 (G) analyzed in Fig. 2C and 2E. Expression in cell lysate was analyzed using 1 OD unit (Ypt7) or 4 OD units (Vps21) loaded onto SDS gel for subsequent western blotting. Protein was visualized using antibodies against Ypt7 and Vps21. Tom40 decoration served as loading control. **(H)** Analysis of autophagic flux. Autophagic flux was analyzed in the indicated strains using the PhoΔ60 assay, for details see methods. (P‑value for *atg9*∆ cells over wild-type after 2h starvation **p<0.01 using two sample t-test). **(I)** Analysis of purified Mon1-Ccz1 complex with mutation of Mon1^L95A,L96A^. GEF complexes were purified as described in the method section and analyzed by SDS-PAGE and Coomassie staining.

**Figure S3. Analysis of the binding interface between Mon1 and the Rab5-like Ypt10 protein.**

(**A**) Carton representation of the *S.c.* Mon1-Ccz1-Ypt7-Ypt10 complex model color-coded according to the pLDDT values. (**B**) Plots of the per-residue local confidence score (on a scale from 0 – 100) and of the predicted aligned error (PAE). (**C**) Carton representation of the modelled complex subunits color-coded according to the pLDDT values. The proposed binding interfaces of Mon1 and Ypt10 are highlighted. (**D**) Close-up of the modeled *S.c.* Mon1-Ypt10 interface (**E**), the modeled *D.m.* Mon1-Rab5 interface, and (**F**) the Fuzzy-Rsg1 interface observed in the cryo-EM structure of CPLANE [Langousis et al., 2022] shown in the same orientation. **(G)** Analysis of the GEF complex containing Mon1^W406A^. GEF complexes were purified using tandem-affinity purification and analyzed by SDS-PAGE and Coomassie staining. (**H**) Expression level of Ypt7 in Figure 3I is not affected by Mon1 mutation. Expression of mNeon‑Ypt7 in cell lysate of wildtype and Mon1^W406^ mutants was analyzed using 1 OD unit cells loaded onto a SDS gel for subsequent western blotting. Protein was visualized using an antibody against Ypt7, Tom40 expression served as loading control.

**Table S1.** Strains used in this study

**Table S2.** Plasmids used in this study

**Table S3**. Kinetic constants of Rab7 GEF complexes.
