## Supplemental Table S1 for "Regulatory sites in the Mon1-Ccz1 complex control Rab5 to Rab7 transition and endosome maturation"

**Table S1.** Yeast strains used in this study

| **Strains** | **Genotype** | **Reference** |
| --- | --- | --- |
| CUY11940 | MATα *leu2-3,112 ura3-52 his3-∆200 trp-∆901 lys2-801 suc2-∆9 GAL ypt7∆::NATNT2 URA3::pRS406-YPT7pr-mNeon-(GGSG)x3-YPT7-YPT7term* | Langemeyer et al., 2020 |
| CUY13443 | MATα *leu2-3,112 ura3-52 his3-∆200 trp-∆901 lys2-801 suc2-∆9 GAL ypt7∆::NATNT2 URA3::pRS406-YPT7pr-mNeon-(GGSG)x3-YPT7-YPT7term MON1∆1-100::KANMX-Mon1pr* | This study |
| CUY13585 | MATα *leu2-3,112 ura3-52 his3-∆200 trp-∆901 lys2-801 suc2-∆9 GAL ypt7∆::NATNT2 URA3::pRS406-YPT7pr-mNeon-(GGSG)x3-YPT7-YPT7term VPS21::hphNT1-Pho5pr-mcherry* | This study |
| CUY13586 | MATα *leu2-3,112 ura3-52 his3-∆200 trp-∆901 lys2-801 suc2-∆9 GAL ypt7∆::NATNT2 URA3::pRS406-YPT7pr-mNeon-(GGSG)x3-YPT7-YPT7term MON1∆1-100::KANMX-Mon1pr VPS21::hphNT1-Pho5pr-mcherry* | This study |
| CUY10489 | MATα *his3Δ1 leu2Δ0 lys2Δ0 ura3Δ0 pho13∆::KAN pho8::PHO8∆60* | Reggiori Laboratory |
| CUY10490 | MATα *his3Δ1 leu2Δ0 lys2Δ0 ura3Δ0 pho13∆::KAN pho8::PHO8∆60 atg9∆::URA* | Reggiori Laboratory |
| CUY13880 | MATα *his3Δ1 leu2Δ0 lys2Δ0 ura3Δ0 pho13∆::KAN pho8::PHO8∆60 MON1∆1-100::natNT2-Mon1pr* | This study |
| CUY12819 | MATa *his3∆200 leu2∆0 met15∆0 trp1∆63 ura3∆0 CCZ1::TRP1-GAL1pr CCZ1::TAP-hphNT1 mon1::kanMX GAL::pRS406-GAL1pr-MON1* | This study |
| CUY13278 | MATa *his3∆200 leu2∆0 met15∆0 trp1∆63 ura3∆0 CCZ1::TRP1-GAL1pr CCZ1::TAP-hphNT1 mon1::kanMX GAL::pRS406-GAL1pr-MON1 L95A L96A* | This study |
| CUY13882 | MATa *his3∆200 leu2∆0 met15∆0 trp1∆63 ura3∆0 CCZ1::TRP1-GAL1pr CCZ1::TAP-hphNT1 mon1::kanMX GAL::pRS406-GAL1-MON1 W406A* | This study |
| SEY6210 | MATα *leu2-3,112 ura3-52 his3-∆200 trp-∆901 lys2-801 suc2-∆9 GAL* | Reggiori Laboratory |
| CUY14281 | MATα *leu2-3,112 ura3-52 his3-∆200 trp-∆901 lys2-801 suc2-∆9 GAL mon1::kanMX LEU::pRS405 Mon1pr-Mon1-Mon1term* | This study |
| CUY14282 | MATα *leu2-3,112 ura3-52 his3-∆200 trp-∆901 lys2-801 suc2-∆9 GAL mon1::kanMX LEU::pRS405 Mon1pr-Mon1 W406A-Mon1term* | This study |
| CUY14283 | MATα *leu2-3,112 ura3-52 his3-∆200 trp-∆901 lys2-801 suc2-∆9 GAL mon1::kanMX LEU::pRS405 Mon1pr-Mon1 W406K-Mon1term* | This study |
| CUY14285 | MATα *leu2-3,112 ura3-52 his3-∆200 trp-∆901 lys2-801 suc2-∆9 GAL mon1::kanMX LEU::pRS405 Mon1pr-Mon1 L95A L96A-Mon1term* | This study |
| CUY13605 | MATα *leu2-3,112 ura3-52 his3-∆200 trp-∆901 lys2-801 suc2-∆9 GAL ypt7∆::NATNT2 URA3::pRS406-YPT7pr-mNeon-(GGSG)x3-YPT7-YPT7term mon1::kanMX LEU::pRS405 Mon1pr-Mon1-Mon1term* | This study |
| CUY14287 | MATα *leu2-3,112 ura3-52 his3-∆200 trp-∆901 lys2-801 suc2-∆9 GAL ypt7∆::NATNT2 URA3::pRS406-YPT7pr-mNeon-(GGSG)x3-YPT7-YPT7term mon1::kanMX LEU::pRS405 Mon1pr-Mon1 W406A-Mon1term* | This study |
| CUY14288 | MATα *leu2-3,112 ura3-52 his3-∆200 trp-∆901 lys2-801 suc2-∆9 GAL ypt7∆::NATNT2 URA3::pRS406-YPT7pr-mNeon-(GGSG)x3-YPT7-YPT7term mon1::kanMX LEU::pRS405 Mon1pr-Mon1 W406K-Mon1term* | This study |
| CUY12706 | MATα *leu2-3,112 ura3-52 his3-∆200 trp-∆901 lys2-801 suc2-∆9 GAL mon1::kanMX* | This study |
