## Supplemental Table S2 for "Regulatory sites in the Mon1-Ccz1 complex control Rab5 to Rab7 transition and endosome maturation"

**Table S2.** Plasmids used in this study

| **Protein** | **Backbone** | **Reference** |
| --- | --- | --- |
| Ypt7 | pET24d-GST-TEV- | Lachmann et al., 2012 |
| Ypt10 | pET24d-GST-TEV- | Lachmann et al., 2012 |
| d.m. Rab5 | pET24d-GST-TEV- | Langemeyer et al., 2020 |
| d.m. Rab7 | pET24d-GST-TEV- | Langemeyer et al., 2020 |
| Bet2-Bet4 | pCDF-DUET*-*1 His-TEV | Thomas et al., 2016 |
| Mrs6 | pET30 | Gift from K.Alexandrov |
| Gdi1 | pGEX-6P | Thomas et al., 2016 |
| d.m. GDI | pET28a-His-Sumo | Langemeyer et al., 2020 |
| GST- PreSc -d.m. Mon1-d.m. Ccz1-3xFlag-d.m. CG8270 | pBig1a- | Langemeyer et al., 2020 |
| GST- PreSc -d.m. Mon1  Δ1-40-d.m.Ccz1-3xFlag- d.m. CG8270 | pBig1a- | This study |
| GST- PreSc -d.m. Mon1  Δ1-50-d.m.Ccz1-3xFlag- d.m. CG8270 | pBig1a- | This study |
| GST- PreSc -d.m. Mon1  Δ1-100-d.m.Ccz1-3xFlag- d.m. CG8270 | pBig1a- | This study |
| GST- PreSc -d.m. Mon1 I47A I48A-d.m.Ccz1-3xFlag- d.m. CG8270 | pBig1a- | This study |
| GST- PreSc -d.m. Mon1 W334A-d.m.Ccz1-3xFlag- d.m. CG8270 | pBig1a- | This study |
| YPT7pr-mNeon-YPT7-YPT7term | pRS406 | Langemeyer et al., 2020 |
| MON1pr-MON1-MON1term | pRS405 | This study |
| MON1pr-MON1 W406A-MON1term | pRS405 | This study |
| MON1pr-MON1 W406K-MON1term | pRS405 | This study |
| MON1pr-MON1 L95A L96A-MON1term | pRS405 | This study |
| GAL1pr-MON1 | pRS406 | This study |
| GAL1pr-MON1 W406A | pRS406 | This study |
| GAL1pr-MON1 L95A L96A | pRS406 | This study |
