## Supplemental Table S3 for "Regulatory sites in the Mon1-Ccz1 complex control Rab5 to Rab7 transition and endosome maturation"

**Table S3. Kinetic constants of Rab7 GEF complexes**

The following tables give an overview on the measured values used to calculate the fold-change in GEF-activity.

Part 1. k_obs_ values (average) for liposome GEF assays using 6.25 nM D.m. Mon1-Ccz1-Bulli. Related to Figure 1E.

| Construct | k_obs_ value  [s^-1^] | STDV |
| --- | --- | --- |
| Trimer wt | 1,39E-03 | 2,94E-05 |
| Trimer Mon1 Δ40 | 2,09E-03 | 1,16E-04 |
| Trimer wt | 1,49E-03 | 3,05E-04 |
| Trimer Mon1 Δ50 | 3,43E-03 | 1,25E-03 |
| Trimer wt | 1,66E-03 | 5,73E-04 |
| Trimer Mon1 Δ100 | 4,68E-03 | 1,25E-03 |
| Trimer wt | 1,87E-03 | 4,49E-04 |
| Trimer Mon1 I47A I48A | 2,99E-03 | 3,77E-04 |

Part 2. K_cat_/k_m_ values (average) for in solution GEF assays using D.m. Mon1-Ccz1-Bulli. Related to Figure 1F.

| Construct | K_cat_/k_m_ value in solution [M^-1^s^-1^] | STDV |
| --- | --- | --- |
| Trimer wt | 1,49E+05 | 1,94E+04 |
| Trimer Mon1 ∆100 | 1,58E+05 | 2,27E+04 |

Part 3. k_obs_ values (average) for liposome GEF assays using 12.5 nM S.c. Mon1-Ccz1. Related to Figure 2I.

| Construct | k_obs_ value [s^-1^] | STDV |
| --- | --- | --- |
| wt | 1,52E-03 | 2,55E-04 |
| Mon1 L95A L96A | 2,06E-03 | 1,93E-04 |

Part 4. K_cat_/k_m_ values (average) for in solution GEF assays using S.c. Mon1-Ccz1. Related to Figure 2I.

| Construct | K_cat_/k_m_ value in solution  [M^-1^s^-1^] | STDV |
| --- | --- | --- |
| wt | 2,45E+04 | 6,98E+03 |
| Mon1 L95A L96A | 2,78E+04 | 3,79E+03 |

Part 5 k_obs_ values (average) for liposome GEF assays using 25 nM S.c. Mon1-Ccz1. Related to Figure 3G.

| Construct | k_obs_ value  [s^-1^] | STDV |
| --- | --- | --- |
| wt | 3,11E-03 | 2,11E-04 |
| Mon1 W406A | 1,80E-03 | 8,41E-05 |

Part 6 K_cat_/k_m_ values (average) for in solution GEF assays using S.c. Mon1-Ccz1. Related to Figure 3G.

| Construct | K_cat_/k_m_ value in solution  [M^-1^s^-1^] | STDV |
| --- | --- | --- |
| wt | 2,46E+04 | 7,11E+03 |
| Mon1 W406A | 3,03E+04 | 7,16E+02 |

Part 7 k_obs_ values (average) for liposome GEF assay using 6.25 nM D.m. Mon1-Ccz1-Bulli. Related to Figure 4C.

| Construct | k_obs_ value  [s^-1^] | STDV |
| --- | --- | --- |
| Trimer wt | 2,25E-03 | 4,80E-04 |
| Trimer Mon1 W334A | 8,62E-04 | 3,28E-04 |

Part 8 K_cat_/k_m_ values (average) for in solution GEF assay using D.m. Mon1-Ccz1-Bulli. Related to Figure 4C.

| Construct | K_cat_/k_m_ value in solution  [M^-1^s^-1^] | STDV |
| --- | --- | --- |
| Trimer wt | 1,14E+05 | 3,83E+04 |
| Trimer Mon1 W334A | 1,22E+05 | 2,48E+04 |
